# scVIP: personalized modeling of single-cell transcriptomes for developmental and disease phenotypes

**DOI:** 10.64898/2026.04.20.717759

**Authors:** Hsin-Yu Lai, Yehchan Yoo, Andreas Tjärnberg, Ruoxin Li, Ziyuan He, Kyle J. Travaglini, Qi Qiao, Anamika Agrawal, Omar Kana, Cindy van Velthoven, Jeff Carroll, Mark Gillespie, Shubhabrata Mukherjee, David W. Fardo, Xiaojun Li, Ed Lein, Mariano Ignacio Gabitto

## Abstract

Single-cell transcriptomics resolves cellular heterogeneity within individuals, but connecting molecular states to individual-level phenotypes requires frameworks that explicitly bridge these scales. We present scVIP, a generative model that links gene expression, cell-type composition, and phenotypic measurements within a single probabilistic model, which enables accurate phenotype prediction and interpretable trajectory inference. A cell-type-aware multi-instance learning architecture learns donor embeddings that capture progression while localizing phenotype-associated signals to specific cell populations. Applied across four settings, scVIP accurately predicts cortical developmental age (Pearson r = 0.95), characterizes Huntington’s disease progression (concordance correlation coefficient = 0.90), integrates two Alzheimer’s disease cohorts recovering disease-relevant microglial and astrocytic programs, and distinguishes healthy from ACPA-positive individuals and non-progressors from early RA individuals, identifying inflammatory T cell programs associated with disease. scVIP enables principled analysis of how cellular states collectively shape organism-level phenotypes across development and disease.

## Introduction

Single-cell genomics has transformed our ability to resolve cellular identities and transcriptional programs at unprecedented resolution [1, 2]. Most computational approaches analyze individual cells as the fundamental unit of observation, enabling the identification of cell types, transcriptional states, and developmental trajectories. However, many biological and clinical questions concern how collections of cells function together within tissues or organisms. Cellular states do not act in isolation: gene expression programs shape cell behavior, cells interact through shared environments and signaling, and shifts in cellular composition collectively determine physiological and pathological outcomes. Understanding these coordinated effects therefore requires moving beyond individual cells to model their collective behavior.

Yet a central challenge remains: linking cellular heterogeneity to phenotypes measured at the level of tissues or individuals. Molecular profiles are routinely measured at single-cell resolution, whereas phenotypic variables such as disease stage, pathological burden, or clinical outcome are typically recorded at the level of donors or samples. Because these phenotypes arise from coordinated activity across multiple cellular populations, pooling cells across individuals can obscure subject-specific signals that are critical for understanding disease mechanisms [3]. Analytical frameworks that fail to account for this hierarchical structure may either collapse meaningful heterogeneity or attempt to predict phenotypes without grounding predictions in the cellular states that generate them. Existing strategies, including differential expression within predefined cell types or sample-level embedding methods [4–6], capture aspects of this variation but do not explicitly model how latent individual states jointly influence gene expression, cell-type composition, and phenotypic measurements.

This limitation is particularly evident across a wide range of biological contexts, from neurodegenerative diseases such as Alzheimer’s disease (AD), where cellular composition and pathological burden vary widely across individuals [7–9], to autoimmune conditions such as rheumatoid arthritis (RA), where systemic immune alterations can precede clinical diagnosis. In both settings, disease progression unfolds along continuous biological trajectories that are only partially captured by discrete clinical assessments such as Braak staging [10–12] or clinical classification criteria. A further challenge is that clinical single-cell studies are rarely conducted in isolation: datasets from independent cohorts often employ different measurement protocols, staging criteria, and phenotypic variables, making direct integration difficult without discarding phenotypic structure.

Recent approaches have begun to address some aspects of these challenges through multi-instance learning architectures or foundation model embeddings [13–16]. However, these methods are predominantly discriminative: they learn mappings from cellular observations to phenotypic labels without explicitly modeling the generative process linking genes, cell-type composition, and individual-level states. As a result, they cannot naturally handle partially observed or inconsistently defined phenotypes across cohorts, do not model compositional shifts jointly with gene expression, and require large labeled cohorts to generalize, which limits their applicability in the small-donor clinical settings where single-cell studies most commonly operate. What is needed is a framework that simultaneously represents all three levels of analysis: genes active within cells, cell types present within a sample, and phenotypic outcomes observed across individuals, while treating missing measurements as partially observed variables and inferring latent individual-level states that jointly explain molecular variation, multicellular organization, and phenotypic outcomes.

Here we present scVIP (single-cell Variational Inference with Personalization), a generative framework that, unlike discriminative approaches, explicitly models the joint distribution over gene expression, cell-type composition, and phenotypic measurements through a unified hierarchical representation spanning genes, cells, and individuals (Fig. 1A). Building on deep generative models for single-cell analysis [17–19], scVIP introduces an explicit latent variable representing each individual’s developmental or disease state. This latent state jointly influences gene expression, cell-type composition, and phenotypic measurements, enabling structured information flow from molecular variation to multicellular organization and ultimately to subject-level outcomes. To infer this representation, scVIP combines variational Bayesian inference with a cell-type–aware multi-instance learning architecture [20], in which attention mechanisms learn how progression manifests across distinct cellular populations before integrating these signals into an individual-level embedding.

**Fig. 1.**
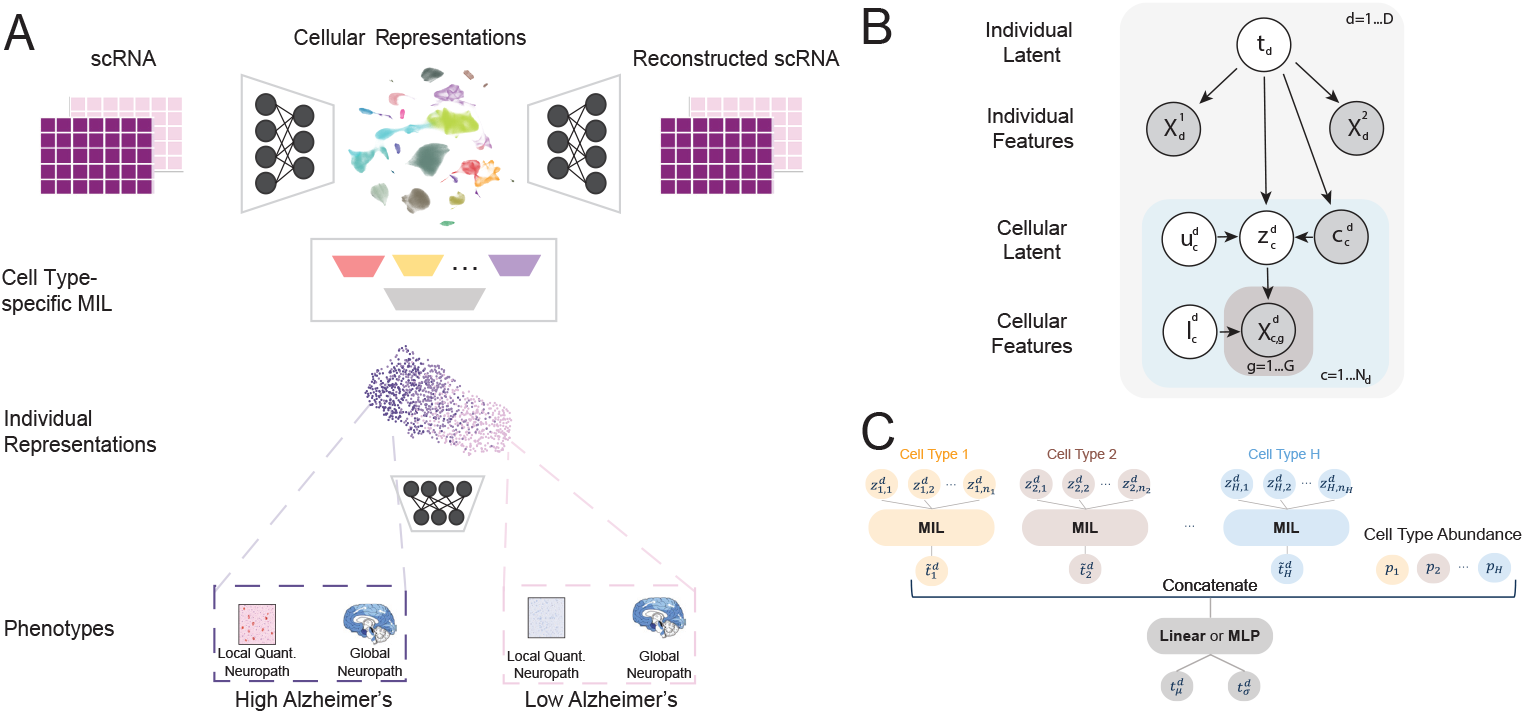
scVIP architecture. (A) Overview of the scVIP framework linking single-cell gene expression, cell-type composition, and phenotypic measurements through a shared individual-level latent representation. (B) Directed acyclic graph for scVIP. (C) Architectures for the cell-type specific MIL (i.e., functions 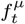 and 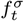 in Eq. (5)). Here, we relabel 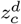 according to the cell type each cell belongs to. We denote *H* as the number of cell types and *n*_*h*_ as the number of cells in cell type *h*. We denote *p*_1_, … , *p*_*H*_ as the cell type abundance of each corresponding cell type. “Concatenate row vectors 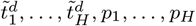 means generating one single row vector: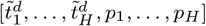. The notation 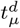 is the output of 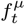, whereas exp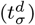 is the output of 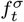 in Eq. (5) in the main manuscript.

We evaluated scVIP across developmental, neurodegenerative, and immune settings and found that the framework consistently learns biologically meaningful and interpretable sample-level representations. The design of scVIP reflects the hierarchical structure of single-cell data and we believe is responsible for its success. First, a deep generative model captures transcriptional variation at the level of individual cells while accounting for technical noise and batch effects. Second, a cell-type–aware multi-instance learning architecture aggregates information across cellular populations within each individual, allowing the model to learn how phenotypic progression manifests differently across cell types. Finally, by modeling cell-type abundances jointly with gene expression, the framework captures compositional shifts that often accompany developmental and disease processes. Together, these components enable scVIP to link molecular variation, multicellular organization, and individual-level phenotypes within a unified probabilistic representation.

## Results

### scVIP recapitulates cortical developmental trajectories

scVIP accurately reconstructs cortical developmental trajectories from single-cell transcriptomes. Cortical development follows a stereotypical progression in which progenitor populations differentiate through intermediate states before reaching mature neuronal and glial identities [21, 22]. To evaluate whether scVIP captures such trajectories, we applied the model to a mouse visual cortex developmental dataset containing 568,674 cells from 53 mice spanning embryonic day 11.5 to postnatal day 56 [23] (see Methods for data preparation and training details). We used developmental age as the phenotype and constrained the scVIP donor embedding to a single latent dimension representing developmental progression. The model was trained on 43 mice and evaluated on a held-out set of 10 animals. scVIP accurately predicted developmental age from single-cell transcriptomes (Pearson *r* = 0.95, mean squared error = 11.08; Fig. 2A–B). Compared with baseline approaches, including a gene-count model, averaged scVI embeddings, and an attention-based multi-instance learning model (see Methods: Comparison with other approaches), scVIP achieved the highest predictive accuracy among the evaluated methods (Fig. 2A), demonstrating that integrating cell-type–specific signals improves phenotype prediction from single-cell data.

**Fig. 2.**
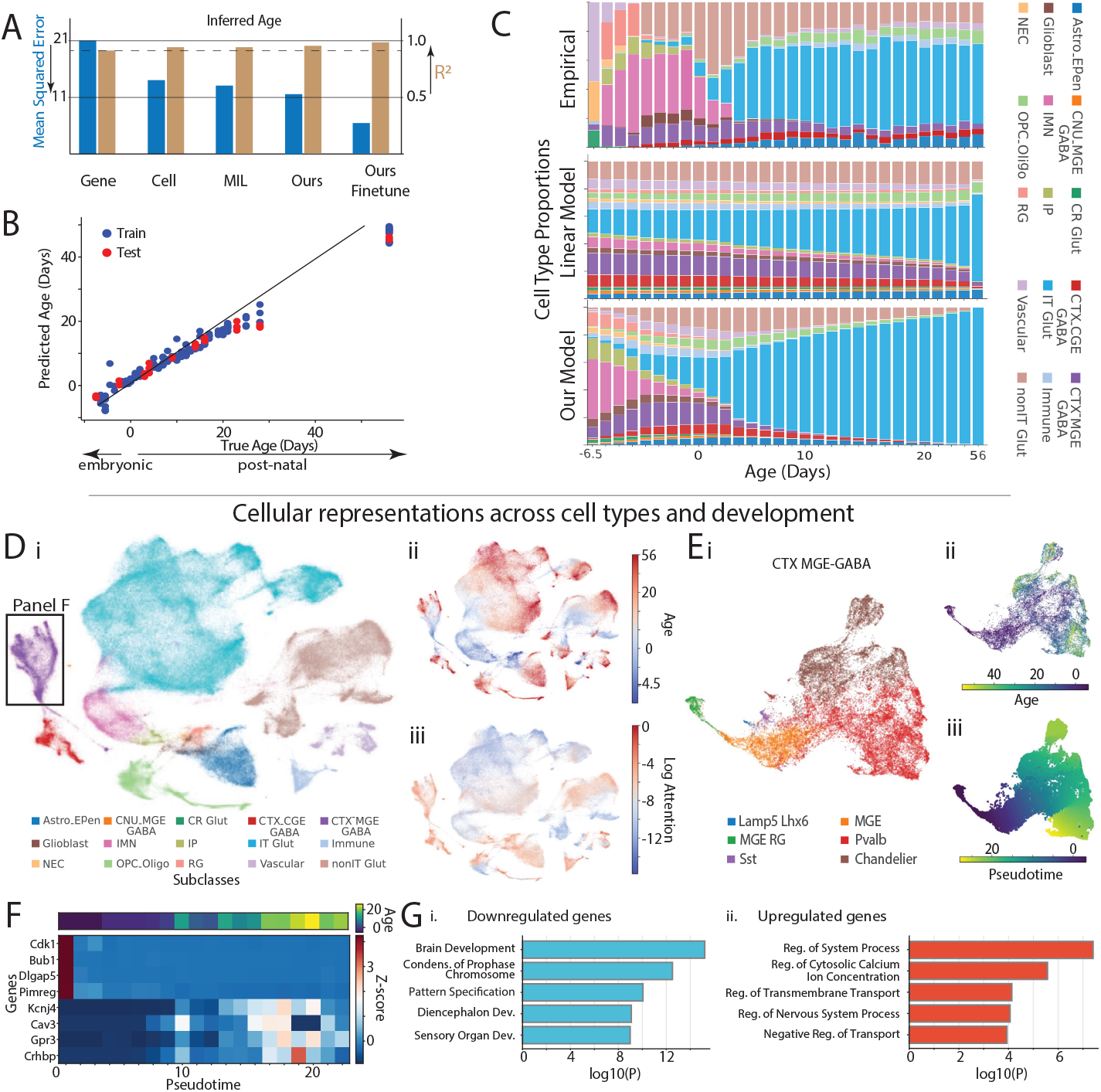
scVIP reconstructs developmental trajectories from single-cell transcriptomes. (A) Performance of scVIP and baseline models in predicting developmental age from single-cell data. Predicted versus observed ages for training and held-out mice. (C) Cell type proportions across age of Top, empirical distribution; Middle, fit of linear model; Bottom, inferred cell type proportions by scVIP. (D) UMAP of cell embeddings colored by i) cell class, ii) developmental age and iii) scVIP attention weights. (E) UMAP of CTX–MGE GABA cells colored by i) subclass, ii) age, and iii) inferred pseudotime. (F) Relationship between developmental age and inferred pseudotime, with representative genes showing differential expression along the trajectory. (G) Gene ontology enrichment of genes differentially expressed along pseudotime.

Beyond prediction, scVIP recovered biologically meaningful developmental structure. Because scVIP explicitly models the dependence between donor-level states and cell-type abundances, we used the mouse visual cortex developmental dataset to evaluate the compositional changes inferred by scVIP. The inferred abundance dynamics closely matched the empirical distributions observed in the dataset (Fig. 2C), correctly capturing early enrichment of progenitor populations such as IMN, IP, Glioblast cells, and the emergence of mature neuronal populations at later stages. More importantly, scVIP captured transient cell populations that a simple linear model fails to recover. Minor overestimation was observed for the most abundant excitatory population (IT Glut) at later developmental stages, likely reflecting the large gap between the final two postnatal time points (postnatal days 22 and 53).

In addition, UMAP visualization of the learned cell embeddings revealed a continuous ordering of cells from early to late developmental stages (Fig. 2D). Within the CTX–MGE GABA lineage, the model reconstructed known interneuron developmental trajectories, including transitions from radial glia to mature Sst and Pvalb interneurons (Fig. 2E), consistent with previously described lineage relationships [23]. To identify transcriptional programs associated with this trajectory, we compared gene expression between highly attended CTX–MGE GABA cell clusters corresponding to early and late developmental stages. Bayesian differential expression (see Methods and Supplementary Data 1) and gene ontology analysis revealed enrichment of cell-cycle and developmental processes among early-stage cells and neuronal signaling pathways among late-stage cells (Fig. 2F–G). Representative genes included *Cdk1, Bub1, Dlgap5*, and *Pimreg* in early cells and *Kcnj4, Cav3, Gpr3*, and *Crhbp* in later stages. Together, these results show that scVIP not only predicts developmental age from single-cell data but also reconstructs developmental trajectories and identifies cell-type–specific molecular programs associated with progression.

### scVIP reveals cell-type-specific signatures of Huntington’s disease (HD) progression

We next evaluated whether scVIP captures transcriptional signals associated with neurodegenerative disease progression, first focusing on Huntington’s disease (HD). HD is caused by CAG repeat expansion in the *HTT* gene, and disease burden is commonly approximated using the CAG-age product (CAP) score [24]. We applied scVIP to single-nucleus RNA-seq data from the anterior caudate nucleus comprising 581,273 cells from 50 individuals with HD and 53 controls [25]. scVIP organized donor embeddings according to CAP score (Fig. 3A) and accurately predicted CAP values in held-out individuals (concordance correlation coefficient = 0.90; Fig. 3B). The model also distinguished HD from control donors with an F1-score of 0.95. Compared with baseline approaches, scVIP achieved higher predictive accuracy, indicating that its hierarchical representation preserves disease-associated variation in single-cell transcriptomes.

**Fig. 3.**
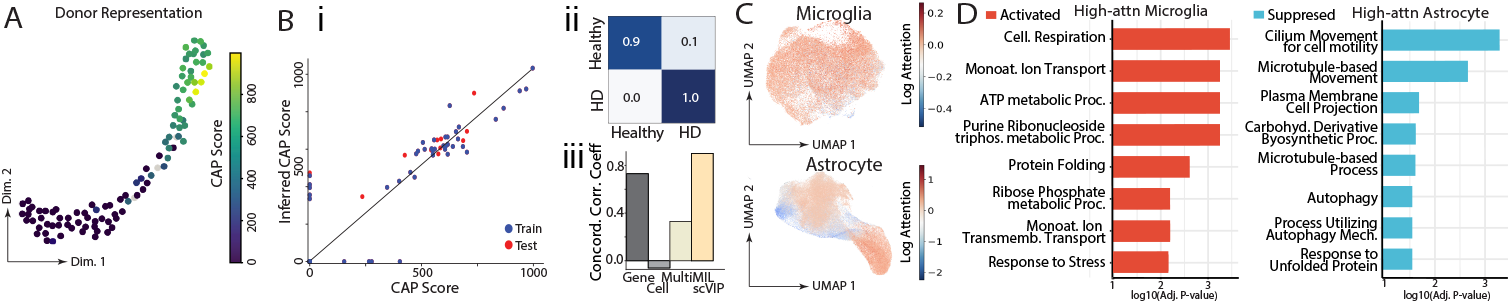
scVIP effectively describes the progression of Huntington’s disease (HD). (A) UMAP of donor embeddings colored by CAP score. (B) Prediction of CAP score and disease status. i) Predicted versus observed CAP scores in the training and test sets. ii) Confusion matrix for control versus HD classification. iii) Performance comparison with baseline methods. (C) UMAPs of microglia and astrocytes colored by attention weights. (D) Gene ontology enrichment of genes differentially expressed between high- and low-attention microglia cells (left) and astrocytes (right).

Attention weights highlighted cellular populations that most strongly contributed to the inferred donor representations (Fig. 3C). Given that neuroinflammation and the loss of glial support are central to HD pathogenesis, we specifically examined microglia and astrocytes, as these cells undergo profound transcriptional remodeling and functional shifts that directly contribute to neuronal vulnerability [26–28]. In microglia, cells receiving higher attention weights (top 20%) showed elevated expression of complement genes (*C1QA, C1QB, C1QC*) and lipid metabolism markers (*APOE, APOC1*), together with increased expression of mitochondrial genes involved in oxidative phosphorylation (see Supplementary Data 2 for the complete result).

Gene ontology enrichment analysis of these cells revealed pathways related to cellular respiration, ion transport, and stress responses (Fig. 3D). These transcriptional signatures are consistent with a metabolically active and stress-responsive microglial state associated with HD progression, consistent with previous reports of complement activation and mitochondrial dysfunction in HD [29–31]. In astrocytes, high-attention cells showed reduced expression of genes related to ciliary structure and microtubule organization, including *CFAP61, CFAP47*, and *DNAH11*. Enrichment analysis revealed suppression of cilium-associated pathways (Fig. 3D), suggesting disruption of astrocytic structural programs during HD progression (see Supplementary Data 2 for the complete result). Although attention weights do not imply causal importance, they provide a useful mechanism for identifying cellular populations that contribute most strongly to the inferred donor-level representation. Together, these results illustrate how scVIP identifies disease-associated cellular states from single-cell transcriptomic data.

### scVIP harmonizes Alzheimer’s disease cohorts and identifies disease-associated cellular programs

We next applied scVIP to analyze single-nucleus RNA-seq datasets from two major Alzheimer’s disease (AD) cohorts (ROSMAP [7] and SEA-AD [8]), and study the possibility of integrating them in a common embedding space. The combined dataset contained 1,034,366 glial cells profiled from dorsolateral prefrontal cortex, prefrontal cortex, and middle temporal gyrus. Because measurement protocols differ across cohorts, we used neuropathological staging variables (Braak stage, Thal phase, CERAD score, and ADNC; see Supplementary Note 2 for explanations and references) to provide consistent supervision while allowing cohort-specific measurements to remain partially observed. scVIP successfully harmonized donors across cohorts, as shown by the integrated donor embeddings (Fig. 4A). Ordering donors along the pseudotime trajectory inferred from these embeddings revealed a monotonic increase in tau pathology measurements, indicating that the model recovered a shared disease progression axis. Across multiple quantitative pathology measures and staging variables, scVIP outperformed baseline methods in predicting neuropathological burden (Fig. 4B) (see Supplementary Note 1 for explanations of metrics).

**Fig. 4.**
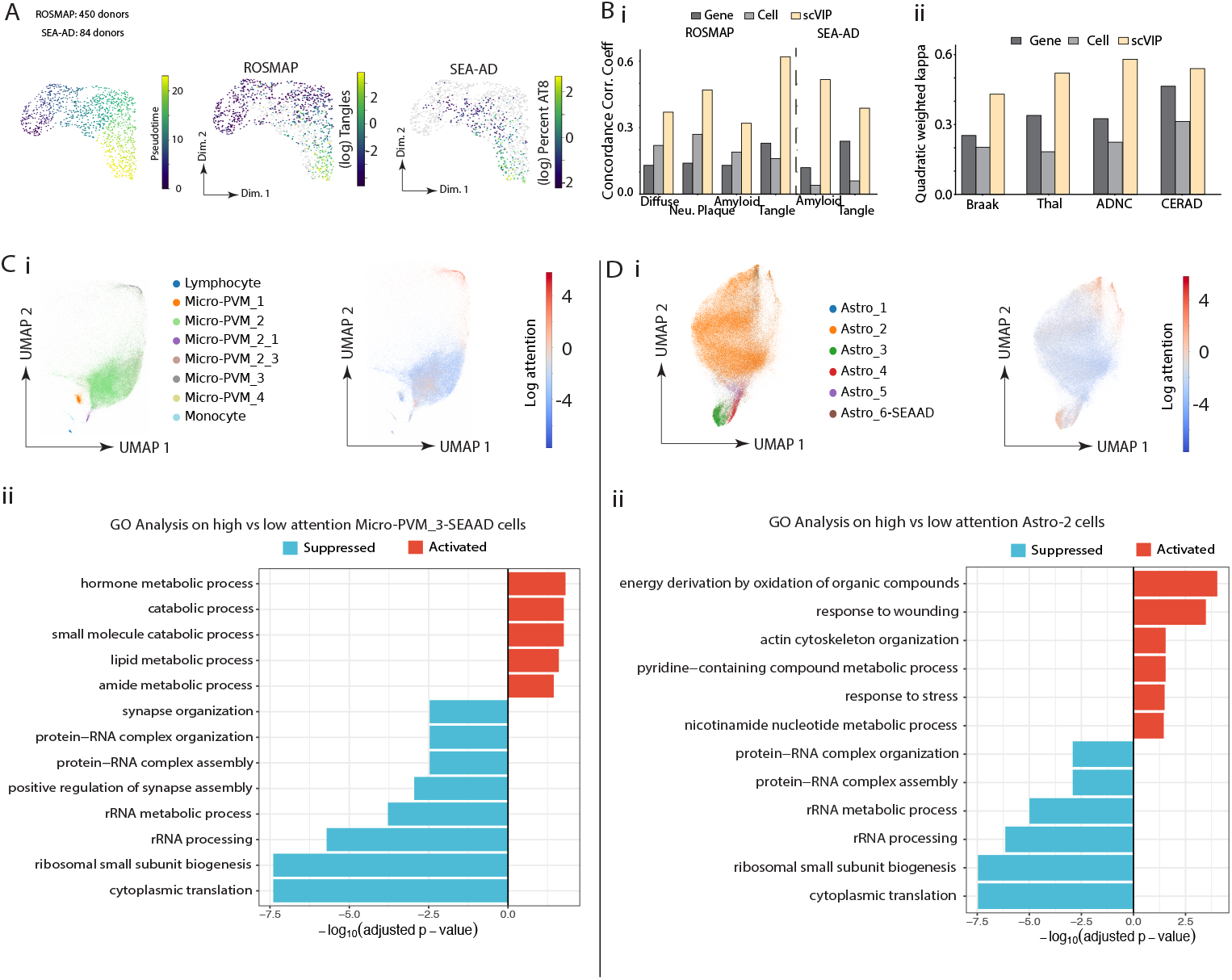
scVIP effectively describes the progression of Alzheimer’s disease (AD). (A) UMAP of donor embeddings showing integration of ROSMAP and SEA-AD cohorts, colored by pseudotime and tau-tangle measurements in each cohort. (B) Prediction performance for i) quantitative pathology measurements and ii) staging. (C) i) UMAP of microglia colored by supertype and attention weights. ii) Gene ontology enrichment of genes associated with high-attention Micro-PVM 3. (D) i) UMAP of astrocyte colored by supertype and attention weights. ii) Gene ontology enrichment of genes associated with high-attention Astro 2.

Attention analysis localized disease-associated signals to specific glial populations. As in the analysis of HD, we further extended our analysis to microglia and astrocytes in AD, as these populations represent the primary cellular drivers of disease progression; while microglial pathways are heavily enriched for AD genetic risk, astrocytes undergo a fundamental loss of homeostatic support and gain of neurotoxic functions that directly correlate with cognitive decline [32]. In microglia, high-attention cells corresponded to the Microglia-PVM3 supertype and showed transcriptional signatures consistent with AD-associated microglia (see Supplementary Data 3 for the full result), including upregulation of lipid metabolism and phagocytic activation genes such as *GPNMB, SPP1*, and *MYO1E*, together with downregulation of homeostatic markers including *CX3CR1* and *P2RY13*. Gene ontology enrichment highlighted lipid metabolic and catabolic processes (Fig. 4C). In astrocytes, high-attention cells corresponded to the Astro-2 supertype and exhibited transcriptional features of reactive astrocytes (Fig. 4D; see Supplementary Data 3 for the full result), including downregulation of translational machinery genes and increased expression of stress-response and extracellular matrix genes such as *CD44* and *SERPINA3*. Enrichment analysis indicated suppression of translational pathways and activation of metabolic and stress-related programs. Notably, these AD-associated signatures differed from those identified in HD, highlighting disease-specific cellular responses captured by scVIP.

### scVIP identifies T cell inflammatory programs in rheumatoid arthritis (RA)

To evaluate whether scVIP captures transcriptional signals associated with autoimmune disease, we applied the framework to peripheral blood mononuclear cells from two cohorts totaling 6,664,701 cells from 185 donors. The ALTRA cohort [33] comprised 89 donors and 2,029,864 cells collected across two study sites, including ACPA-negative healthy controls, ACPA-positive at-risk individuals who either remained non-progressors (NONC) or subsequently progressed to clinical RA (CONV), and ACPA-positive early RA individuals (ERA). The SoundLife cohort [34] contributed 96 healthy control donors totaling 4,634,837 cells after stratification to improve data balance across cohorts; see Methods for details on data preparation. Because all NONC, CONV, and ERA individuals in this cohort were ACPA-positive, we trained scVIP jointly on two RA-related classification tasks: distinguishing ACPA-negative healthy controls from ACPA-positive individuals, and distinguishing NONC from ERA disease states. Labels not used for a given training objective were treated as unknown in a semi-supervised framework. To prevent leakage, train/evaluation splits were performed by donor: all bags from a given subject were assigned exclusively to either training or evaluation.

scVIP organized donor embeddings along axes that separated disease groups without obvious confounding by age or biological sex (Fig. 5A). We evaluated classification performance at the bag level using macro-averaged F1 to account for class imbalance, and compared scVIP with gene-count, cell-embedding, and PULSAR baselines. For these baselines, random forest classifiers were trained on fixed feature representations using the same train/evaluation splits. PULSAR was included as a PBMC-focused foundation-model baseline using donor-level embeddings. For the RA analysis, scVIP used a two-dimensional individual-level latent representation to represent the two supervised classification objectives, whereas the baseline classifiers operated on substantially higher-dimensional fixed feature spaces.

**Fig. 5.**
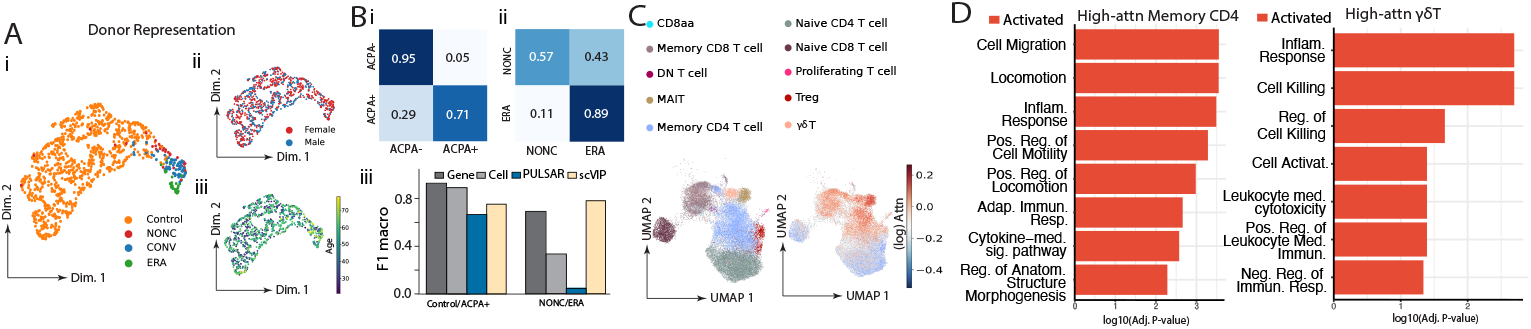
scVIP identifies T cell inflammatory programs in rheumatoid arthritis (RA). UMAP of donor embeddings colored by i) disease group, showing separation of healthy controls, ACPA-positive at-risk individuals, including non-progressors (NONC) and progressors (CONV), and early RA (ERA) donors, ii) biological sex, and iii) age at draw, demonstrating that the learned representation is not confounded by demographic variables. (B) Prediction performance for i) ACPA-negative healthy controls versus ACPA-positive individuals and ii) NONC versus ERA classification, with confusion matrices showing per-class accuracy and bar plots showing macro-averaged F1 for scVIP and baseline approaches, including gene-count, cell-embedding, and PULSAR baselines. (C) UMAP of T cells colored by subtype (left) and attention weights (right), highlighting memory CD4 T cells and *γδ* T cells as the most strongly attended populations. (D) Gene ontology enrichment of genes differentially expressed between high- and low-attention memory CD4 T cells (left) and *γδ* T cells (right).

For the broad ACPA-negative HC/ACPA-positive classification, gene-count and cell-embedding baselines achieved the highest macro-F1 scores (0.89 and 0.86, respectively), while scVIP and PULSAR achieved macro-F1 scores of 0.72 and 0.64 (Fig. 5B). This indicates that the broad distinction between healthy controls and ACPA-positive individuals can be captured by aggregate transcriptional features. For the NON-C/ERA task, which distinguishes ACPA-positive individuals with different clinical disease states, the limited number of ERA donors required splitting each ERA sample into three independent sub-bags for both training and evaluation. scVIP achieved the highest macro-F1 score on this task (0.80), outperforming gene-count, cell-embedding, and PULSAR baselines (0.69, 0.33, and 0.05, respectively; Fig. 5B). Specifically, scVIP correctly classified 57% of NONC bags and 89% of ERA sub-bags, indicating that the scVIP representation was most beneficial for resolving finer disease-state differences within ACPA-positive individuals.

As an exploratory analysis, we further examined model predictions for CONV donors, i.e., ACPA-positive at-risk individuals who subsequently progressed to clinical RA, across longitudinal time points before diagnosis. Most CONV samples were predicted as ERA-like even before clinical diagnosis, suggesting that scVIP may detect circulating immune states shared with early RA prior to clinical diagnosis. These results should be interpreted cautiously given the small number of CONV subjects (n = 16; Supplementary Figure 1).

Attention analysis highlighted memory CD4 T cells and *γδ* T cells among the populations contributing strongly to the inferred donor representations (Fig. 5C). In memory CD4 T cells, high-attention cells showed elevated expression of chemokine receptors and inflammatory or activation-associated genes, including *CCR2, CXCR6, CD70*, and *IL18RAP*, together with tissue-residency, adhesion, and effector-associated markers such as *HOPX* and *LGALS3* (see Supplementary Data 4 for the full result). Gene ontology enrichment analysis revealed pathways related to cytokine-mediated signaling, adaptive immune response, inflammation, locomotion, and cell migration (Fig. 5D), consistent with the established involvement of memory CD4 T cells in RA-associated immune activation. In *γδ* T cells, high-attention cells showed elevated expression of chemokine receptors *CCR1* and *CCR2, IL18RAP*, and immune-signaling or effector-associated genes including *BLK* and *KLRB1* (see Supplementary Data 5 for the full result). Enrichment analysis revealed pathways related to leukocyte-mediated cytotoxicity, cell killing, cell activation, and inflammatory response (Fig. 5D), consistent with cytotoxic and inflammatory programs implicated in rheumatoid arthritis and related autoimmune conditions. Together, these results suggest that scVIP identifies coordinated inflammatory, migratory, and cytotoxic T cell programs in peripheral blood associated with the RA risk/disease spectrum, including immune states that may be detectable before clinical diagnosis.

## Discussion

scVIP introduces a hierarchical generative framework in which a directed acyclic graph explicitly encodes biological relationships from genes to cells to individuals. This design distinguishes scVIP from approaches that either analyze cells in isolation or aggregate them prior to modeling: by coupling cellular and individual levels through a shared latent variable, the framework allows phenotypic supervision to propagate back to cell-level representations without requiring all cells to contribute equally. The cell-type-aware attention mechanism is central to this: rather than treating a sample as an unordered bag of cells, it learns which populations and transcriptional states are most informative for a given phenotype, providing interpretability alongside prediction. This design also makes scVIP naturally suited to cross-cohort harmonization, where phenotypic measurements are partially observed or defined inconsistently across studies—a pervasive challenge in clinical genomics that existing sample-level embedding methods do not directly address.

A practical challenge in evaluating models at this scale is that closely related methods often differ substantially in their supervision, domain assumptions, and accessibility. We therefore selected baselines that tested distinct alternative explanations for scVIP performance, including donor-level gene-expression summaries, aggregated cell embeddings, and multi-instance learning approaches. Recent patient- or sample-level single-cell methods provide important related work, but many are not directly comparable across the settings considered here because they are restricted primarily to categorical prediction [13], require pretrained foundation-model weights that are not publicly available or request-only [14], depend on domain-specific pretrained models [15], or do not naturally support the semi-supervised setting with partially observed phenotypes used by scVIP [16]. In the RA analysis, we additionally compared against PULSAR, a PBMC-focused foundation model, because it provides donor-level immune embeddings in the relevant tissue domain. PULSAR captured some signal for the broad ACPA-negative healthy-control versus ACPA-positive contrast, but performed poorly on the finer NONC versus ERA classification, highlighting that domain-specific foundation-model embeddings do not necessarily resolve cohort-specific disease-state distinctions without task-specific adaptation. In contrast, scVIP learns individual-level representations directly from the target dataset and does not require a large pretrained foundation model matched to the biological domain.

While the model achieves high accuracy in phenotypic prediction and reveals biologically meaningful transcriptional programs, several opportunities remain to further extend the framework. Because scVIP identifies signals that are predictive of target phenotypes, it may highlight only a subset of the molecular processes underlying developmental or disease trajectories. Direct analysis of gene expression dynamics along the learned pseudotime trajectories may uncover additional programs beyond those captured by the supervised objective alone. At the same time, signals correlated with disease progression, including potential technical confounders, may also be captured, motivating future work to more explicitly disentangle biological from nuisance variation. Incorporating additional molecular modalities such as chromatin accessibility or proteomic measurements could provide a more complete view of regulatory processes; because scVIP is modular by design, such extensions are straightforward; for example, the scVI backbone could be replaced with multimodal variants such as MultiVI [19] without altering the overall framework. Finally, incorporating biological priors such as gene regulatory networks or cell–cell communication signals may further improve interpretability and facilitate more precise identification of disease mechanisms.

In the RA application, scVIP was trained jointly on two classification tasks— distinguishing ACPA-negative healthy controls from ACPA-positive individuals and distinguishing NONC from ERA donors—using a shared individual-level representation. This multi-task design enables efficient use of partially labeled data across cohorts, and was most beneficial for the finer NONC/ERA classification, where scVIP outperformed all evaluated baselines. However, the same design also introduces an interpretability limitation: because attention weights are learned from a shared representation, the transcriptional programs identified may reflect signals relevant to both classification axes simultaneously rather than being uniquely attributable to one phenotypic contrast. This limitation is particularly relevant in light of the RA benchmark results, where simple aggregate features performed strongly on the broad HC/ACPA-positive task, whereas scVIP was most beneficial for resolving the more specific NONC/ERA disease-state distinction. A natural extension would be to introduce task-specific attention heads within the multi-instance learning architecture, allowing each classification objective to learn its own cell-type weighting while still sharing the generative backbone. Such a multi-head extension could improve both predictive flexibility and biological interpretation in multi-task disease settings. Additionally, the limited number of ERA and CONV donors means that the preclinical prediction results should be viewed as hypothesis-generating rather than definitive.

A complementary direction is to apply scVIP across a broader range of diseases. As illustrated by the HD, AD, and RA analyses, the model identifies distinct transcriptional programs in each setting, suggesting partially overlapping but disease-specific molecular mechanisms. In neurodegeneration, attention highlighted glial populations: microglia and astrocytes, consistent with their central roles in HD and AD pathogenesis. In autoimmune disease, attention shifted to T cell compartments, identifying effector memory CD4 T cells and *γδ* T cells as the most disease-informative populations, with transcriptional signatures of inflammation, cytokine signaling, and leukocyte migration. Systematically comparing these programs across diseases could reveal pathways shared across conditions as well as those that are disease-specific. Realizing this at scale may benefit from more scalable cell-level encoders, including foundation-model architectures for single-cell data. In principle, the hierarchical DAG underlying scVIP is compatible with such encoders: larger pretrained or foundation-like architectures could replace the current scVI backbone if they are accessible for training or fine-tuning with the appropriate generative and phenotype-supervised objectives. However, fixed embeddings from pretrained models cannot fully substitute for this integration, because they do not allow phenotype- and individual-level losses to shape the cell-level representation within the DAG. Thus, scVIP provides a flexible framework for future extensions using larger encoders, while also remaining applicable in settings where only target-cohort data and standard single-cell generative models are available.

More broadly, scVIP represents a step toward a general framework for linking single-cell molecular measurements to organism-level phenotypes across diverse biological contexts. As single-cell atlases continue to grow in scale and phenotypic depth, models that explicitly bridge cellular and individual-level variation will be essential for understanding how molecular heterogeneity shapes disease risk, progression, and outcome. We anticipate that the principles underlying scVIP—hierarchical generative modeling, personalization, and cell-type-aware design—will generalize beyond neurodegeneration and immune diseases to a wide range of complex diseases where connecting cellular states to clinical phenotypes remains a central challenge.

## Methods

### scVIP model and inference

scVIP extends deep generative models for single-cell transcriptomics by introducing an explicit individual-level latent representation that links cellular measurements to sample-level phenotypes. The framework builds on variational autoencoder (VAE)-based models for gene expression [18, 19], while incorporating a structured mechanism to infer subject-specific developmental or disease states (see Fig. 1B for the directed acyclic graph (DAG)).

Given single-cell gene expression measurements 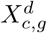 from gene *g* in cell *c* in individual *d*, along with observed phenotypic markers *X*_*d*_, scVIP models the data through a hierarchical latent structure. At the individual level, each sample is associated with a latent variable *t*_*d*_, representing each individual’s underlying developmental or disease state. At the cellular level, each cell is assigned a latent embedding 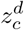 that captures transcriptional variation conditional on both cell-type identity and the individual state as well as its cell type *c*_*c*_.

In the generative component, the individual latent state jointly influences gene expression, cell-type composition, and phenotypic measurements. To be more precise:

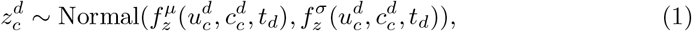

and the cell-type counts conditional on *t*_*d*_ follow the scCODA model [35] with a simplification on the prior for *β*:

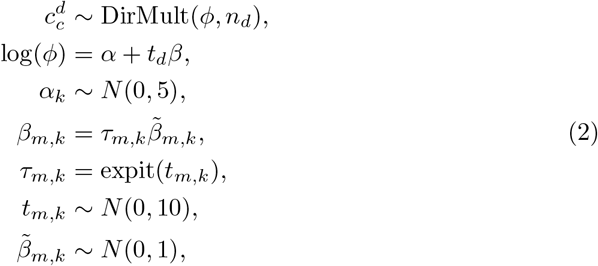

where *n*_*d*_ is the total number of cells in individual *d*. Additionally, we assume the following priors and phenotype likelihoods:

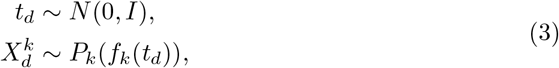

where *f*_*k*_ is defined by linear or MLP layers and *P*_*k*_ is determined based on the data type of 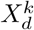 and can be customized: multinomial logistic regression for categorical variables, CORN [36] for ordinal variables, Gaussian for continuous variables, Hurdle Normal [37, 38] for zero-inflated variables. Gene expression is modeled using a negative binomial likelihood, as in scVI [17], with parameters conditioned on both *z*_*c*_ and *t*_*d*_.

For inference, we approximate:

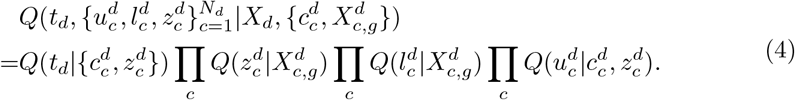

The implementations of 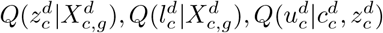 follow standard approaches as in scANVI [39]. To implement

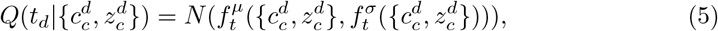

scVIP aggregates information across cells using a multi-instance learning (MIL) architecture [20] with cell-type awareness as described below:

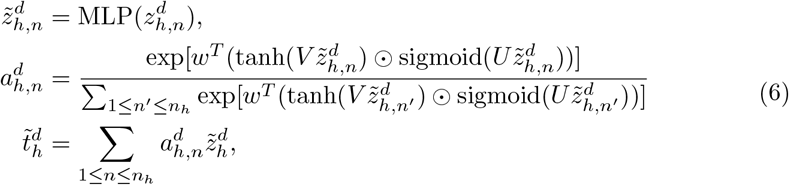

where *h* denotes the cell type each cell belongs to, *n*_*h*_ denotes the number of cells in cell type *h*. The parameters *w, V, U* and the MLP parameters are learned during training. Rather than considering all cells within a sample, scVIP partitions cells by annotated cell type and applies cell-type-specific MIL attention mechanisms to learn how donor state manifests differently across cellular populations — a design in which each attention head operates over a distinct cell-type subpopulation before aggregation, drawing conceptual parallels to multi-head attention in transformer architectures. As shown in Figure 1C, each 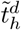 for each cell type as well as cell type abundance are then concatenated and operated through a linear mapping or MLP layers to get the mean 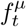 and variance 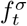 of the Gaussian distribution that defines 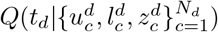).

Training of scVIP follows a two-step procedure. First, we train scVI to obtain initial cell embeddings. When multiple datasets are available and phenotypic measurements may be missing in some datasets (for example, across training and testing sets or across cohorts), we recommend learning cell embeddings using cells from all datasets at this stage. After marking missing phenotypes as NA, scVIP is applied to jointly learn individual-level representations while refining cell embeddings (see Supplementary Figure 2 for visualization of this two-step training). This procedure aims to maximize an evidence lower bound (ELBO), incorporating reconstruction terms and Kullback-Leibler regularization [40] (see Supplementary Note 2 for the loss function). To reduce overfitting, we apply a dropout rate of 0.2 when learning the mapping from the MIL outputs and cell-type abundances to the mean and variance of the individual-level embeddings.

To evaluate differential gene expression between two sets of cells, we adopted the Bayesian differential expression framework implemented in scVI [17]. In this approach, posterior samples of gene expression are obtained from the variational distribution conditioned on each cell’s observed gene counts, batch annotation, and phenotypic information. Because our variational approximation assumes that the latent cell representation 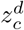 is independent of the phenotypes 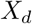 given the observed gene expression 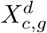, posterior sampling can be implemented in the same manner as in scVI. Assuming cells are independent given the latent variables, these posterior samples are used to estimate the probability that a gene is more highly expressed in one set of cells than in the other. Genes are then ranked according to this posterior probability of differential expression. For developmental data, all genes with posterior probability equaling one were considered. For the rest of the data, except for HD microglia, all genes used in scVIP were ranked using the Bayesian differential expression probability multiplied by the sign of the estimated expression difference. This signed ranking statistic was used as input for gene set enrichment analysis. For HD microglia, ribosomal protein genes were excluded from the ranked gene list before GO enrichment analysis to reduce enrichment of generic translation- and ribosome-associated terms.

### Data preparation

For the developmental dataset, quality control (QC) was performed as described in [23]. For the HD and immune datasets, we selected the top 5,000 highly variable genes. After training scVI, we performed additional QC based on subclass marker genes. Specifically, Leiden clustering was applied to the scVI embeddings, and the distribution of marker gene expression was examined across clusters. Clusters exhibiting elevated expression of marker genes from undesired subclasses and reduced expression of marker genes from the target subclass were identified and removed (thresholds were decided through visualization of the histograms of the cluster expressions and visualization of the cluster locations in the UMAP). For the ROSMAP and SEA-AD datasets, QC was performed following the procedure described in [8]. We then selected 4,351 genes associated with cell types or with potential links to Alzheimer’s disease progression [8].

For the immune data, we merged the blood mononuclear cells (PBMCs) from two cohorts: the ALTRA cohort, a longitudinal study of RA progression, and the SoundLife cohort, comprising healthy controls. The quality-controlled data of both cohorts are accessible [33, 34]. Since SoundLife cohort contains significantly more cells than the ALTRA cohort, we subsampled each donor from the SoundLife cohort to at most have 50,000 cells by stratifying at the supertype level to maintain cell type composition.

### Training details for the datasets

In the developmental dataset, the donor–library pair was used as the batch key. In the HD dataset, library preparation was used as the batch key, and explicitly included sex as a biological covariate. This adjustment was necessary because initial embeddings generated without this covariate exhibited sex-specific clustering within shared cell types, indicating that sexual dimorphism was a dominant driver of transcriptional variance that masked disease-relevant signals. In the AD datasets, cohort was used as the batch key, whereas library preparation was included as a covariate. In the immune datasets, cohort was used as the batch key, whereas library preparation and sex were included as covariates. Model hyperparameters were consistent across all experiments, including four encoder and four decoder layers, 30-dimensional cell embeddings, and a negative binomial gene likelihood.

To facilitate prediction of pathology (as neural network weights are typically initialized near zero), CAP values in the HD dataset were scaled by dividing by 200. For the AD datasets, neuropathological measurements were log-transformed (log(*x* +0.01) for ROSMAP and log(*x* + 0.001) for SEA-AD) to place measurements from the two cohorts on comparable scales.

Training was performed for 100 epochs on the developmental dataset, 20 epochs on the HD dataset, 30 epochs on the combined ROSMAP and SEA-AD dataset, and 20 epochs on the combined Soundlife and ALTRA dataset, all with a 0.001 learning rate. To account for potential differences in staging measurements between ROSMAP and SEA-AD, an additional interaction bias term was included for predicting staging values.

Immune data contained substantial class imbalance and longitudinal sampling, requiring a training design that preserved subject-level independence while providing sufficient bags for model fitting. We defined a bag as a unique combination of subject ID, visit, and batch ID. To prevent leakage, train/evaluation splits were performed at the donor level: all bags from a given subject were assigned exclusively to either training or evaluation. We then applied task-specific bag construction to address class imbalance. For healthy controls, which were substantially overrepresented because of the large SoundLife cohort, only the largest bag from each HC training donor was retained for model fitting. Conversely, because the number of ERA donors was limited, each ERA bag was split into three sub-bags by randomly assigning cells to one of three groups; the same sub-bagging procedure was applied to ERA samples in both training and evaluation.

For the ACPA-negative healthy control (HC) versus ACPA-positive classification task, this procedure yielded 101 HC donors represented by 101 HC training bags and 46 ACPA-positive donors represented by 103 ACPA-positive training bags. The held-out evaluation set contained 26 HC donors represented by 185 HC bags and 12 ACPA-positive donors represented by 27 ACPA-positive bags.

For the NONC versus ERA classification task, this procedure yielded 24 NONC and 24 ERA training bags. The held-out evaluation set contained 7 NONC donors represented by 7 bags and 3 ERA donors represented by 9 sub-bags, and 16 CONV bags for exploratory prediction. HC and CONV donors were treated as unknown for the NONC/ERA objective during training. The model was trained jointly on both classification tasks using a semi-supervised framework, with labels not used for a given training objective marked as unknown.

### Comparison with other approaches

To construct a model based on gene counts, we first reduced gene dimensionality to mitigate overfitting. We applied the local spatial pattern (LSP) score [41], which identifies genes whose expression is locally enriched among neighboring cells on the k-nearest-neighbor graph, where the neighbor graph is defined using the scVI latent space. Gene counts were then normalized at the single-cell level. For each individual (a mouse or donor), we computed the average expression of each gene across all cells. These aggregated gene expression profiles were used to train a random forest model. To construct a model based on cell embeddings, we computed the average of the scVI-derived cell embeddings for each cell type. If there are *H* cell types, this yields *H* averaged embeddings per individual. Cell-type abundances were also included as additional features. These features were concatenated and used to train a random forest model for phenotype prediction.

MultiMIL implementation is available online [5]. In the mouse visual cortex analysis, the multiVAE component (used for learning cell embeddings) was trained jointly with the MIL layer (used for predicting phenotypes from the cell embeddings). However, due to recent changes in the multiMIL codebase, only the MIL layers are available in the current GitHub main branch for application to the HD dataset. To ensure consistency with the latest implementation, we replaced the original multiVAE-based embeddings with cell embeddings learned using scVI, and applied the MIL layers to predict CAP scores.

PULSAR PBMC model weights are publicly available [15]. We applied PULSAR to the immune dataset by representing each 1,024-cell donor-derived bag as a 512-dimensional embedding. For downstream classification, we averaged embeddings across all bags from the same donor and used the resulting 512-dimensional donor-level representation as input features. Random forest classifiers were then trained on these donor-level PULSAR features using the same donor-level train/evaluation split.

In the developmental dataset, we noticed that although scVIP predictions were strongly correlated with ground-truth ages, small systematic biases remained (Fig. 2B). To examine whether these biases reflected limitations of the regression layer rather than the learned representation, we trained a linear model using the inferred donor embeddings *t*_*d*_ as input (denoted as “Ours Finetune” in Fig. 2B). This additional step further improved prediction accuracy, indicating that the latent representation learned by scVIP captures the developmental signal effectively and that full VAE optimization may slightly trade predictive calibration for improved representation generality.

## Data availability

The mouse visual cortex developmental dataset is available from Gao et al. [23]. The Huntington’s disease dataset is available from Handsaker et al. [25]. The ROSMAP single-nucleus RNA-seq dataset analyzed in this study is available through the Synapse repository (accession syn31512863) and was originally described by Green et al. [7]. The SEA-AD MTG and DFC datasets are available through the Seattle Alzheimer’s Disease Brain Cell Atlas (sea-ad.org) and were described by Gabitto et al. [8]. The ALTRA cohort and the SoundLife cohort are both available [33, 34].

## Code availability

We are now providing jupyter notebook files without datasets for reference code to generate figures and also, a capsule with a tutorial that runs on a test data set at https://codeocean.allenneuraldynamics.org/capsule/4344243/tree. In the future, the code is planned to be made available on the scvi-tools, with proper documentation, and tutorials.

## Acknowledgments

This research is sponsored by NIH Grant R01AG082730.

## Author contributions

H.-Y.L. conceptualized and designed the model under M.I.G.’s supervision. H.-Y.L. led the implementation and conducted all experiments on the presented datasets.

Y.Y. refactored the codebase and prepared the reproducible version. A.T., R.L., Z.H., K.J.T., Q.Q., A.A., and O.K. contributed to model development or data cleaning through feedback during exploration. C.V.V., J.C., M.G., S.M., D.W.F., X.L., and E.L. provided domain expertise and feedback on biological interpretation of results. H.-Y.L. and M.I.G. wrote the manuscript with input from all authors. M.I.G. acquired funding and supervised the project.

