## Supplementary figures and texts for "scVIP: personalized modeling of single-cell transcriptomes for developmental and disease phenotypes"

### 1 Supplementary Figure

#### 1.1 Supplementary Figure 1: scVIP predictions on CONV donors at each visit

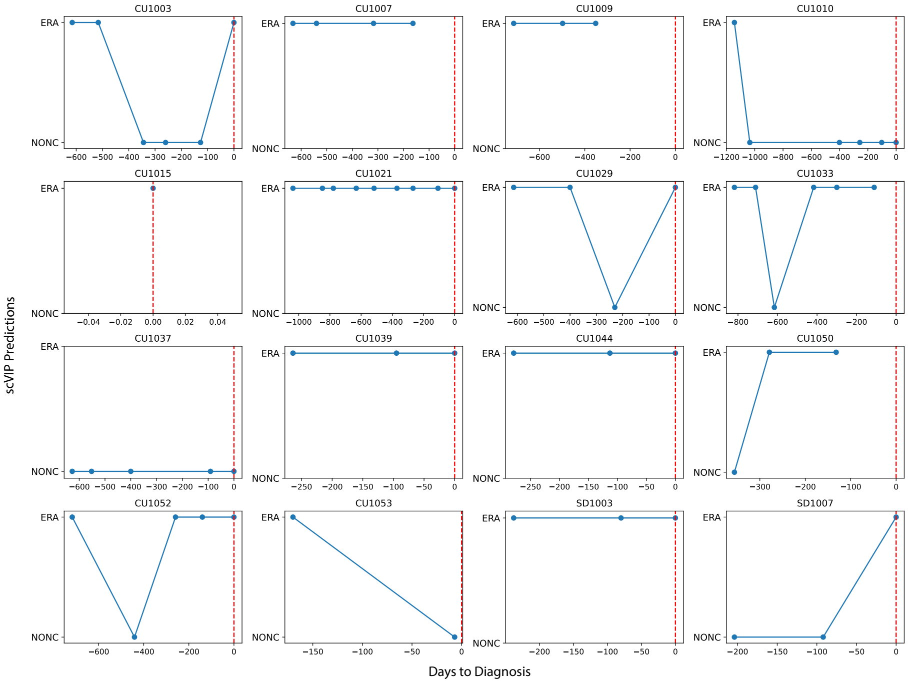

Supplementary Figure 1: **scVIP predictions on CONV donors across longitudinal timepoints.** Each panel shows the binary scVIP NONC/ERA prediction for a single CONV donor (n=16) plotted against days to diagnosis. The red dashed line indicates the day of diagnosis (day 0). The majority of CONV donors are predicted as ERA-like across multiple timepoints, including years before clinical diagnosis. A subset of donors shows variable predictions over time. These results should be interpreted cautiously given the small sample size and binary nature of the predictions.

#### 1.2 Supplementary Figure 2: Training procedures for scVIP

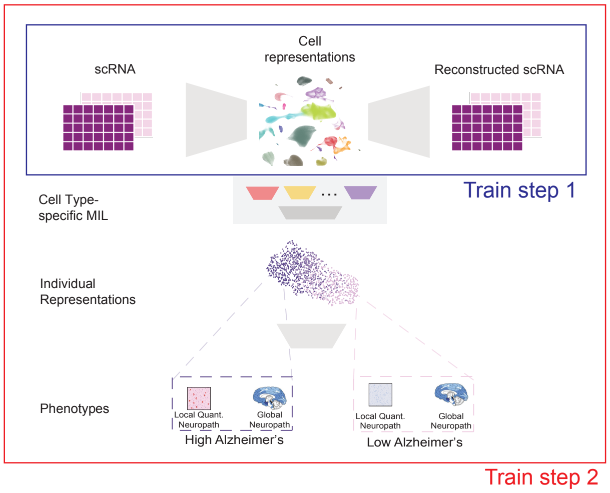

Supplementary Figure 2: Training procedures for scVIP

### 2 Supplementary Notes

#### 2.1 Supplementary Note 1: Explanation of terms

- Quantitative pathology measurements from the AD ROSMAP dataset (see [1] and [2] for more details):
  - Diffuse plaque density: counts of diffuse plaque by region determined by microscopic examination of silver-stained slides
  - Neuritic plaque density: counts of neuritic plaque by region determined by microscopic examination of silver-stained slides
  - Amyloid density: percent area of cortex occupied by amyloid beta where amyloid beta was identified using immunohistochemistry and quantified by image analysis
  - Neurofibrillary tangle (NFT) density: cortical density (per mm2) of NFTs using systematic sampling where NFTs were identified using immunohistochemistry and quantified by image analysis
- Quantitative pathology measurements from the SEA-AD dataset (see [3] for more details):
  - Percent 6e10 positive area: Percent of voxels that stained positive with the 6e10 antibody used to detect amyloid beta peptides, which accumulate throughout the progression of AD.

  - Percent AT8 positive area: Percent of voxels that stained positive with the AT8 antibody used to detect hyperphosphorylated tau, which accumulate throughout the progression of AD.

- AD staging:
  - Braak: Ordinal extent of the anatomical distribution of neurofibrillary tangles (NFTs) [4]
  - Thal: Ordinal extent of the anatomical distribution of amyloid beta plaque deposits [5]
  - CERAD: Ordinal semiquantitative neuritic plaque density [6]
  - ADNC: Ordinal Overall Alzheimer's disease neuropathologic change score [7]

- Performance metrics:
  - Concordance correlation coefficients (CCC): a measure of agreement between two continuous variables [8]. It accounts for both systematic bias and the strength of correlation between the variables.
  - Quadratic weighted kappa (QWK): a variant of Cohen's kappa that applies quadratic weights to account for the ordinality of the data [9]. It is a common measure of agreement between two raters, with the quadratic weights penalizing larger disagreements more heavily than smaller ones.

#### 2.2 Supplementary Note 2: Loss functions

Applying the evidence lower bound (ELBO) for variational inference [10] to our model gives:

$$
\begin{aligned}
&\arg\min -\mathbb{E}_Q[\log P(X_d, X^d_{c,g}, t_d, \{u^d_c, l^d_c, z^d_c, c^d_c\})] \\
&\qquad + \mathbb{E}_Q[\log Q(\{u^d_c, l^d_c, z^d_c\}, t_d | X_d, \{X^d_{c,g}, c^d_c\})] \\
&= \arg\min -\mathbb{E}_Q[\log P(t_d) + \log P(X_d|t_d) + \log P(u^d_c) \\
&\qquad + \log P(c^d_c|t_d) + \log P(z^d_c|t_d, c^d_c, u^d_c) \\
&\qquad + \log P(l^d_c) + \log P(X^d_{c,g}|l^d_c, z^d_c)] \\
&\qquad + \mathbb{E}_Q[\log Q(z^d_c|X^d_{c,g}) + \log Q(l^d_c|X^d_{c,g}) \\
&\qquad + \log Q(t_d|\{c^d_c, z^d_c\}) + \log Q(u^d_c|c^d_c, z^d_c)] \qquad (1) \\
&= \arg\min \mathrm{KL}(Q(z^d_c|X^d_{c,g})||P(z^d_c|t_d, c^d_c, u^d_c)) \\
&\qquad + \mathrm{KL}(Q(l^d_c|X^d_{c,g})||P(l^d_c)) \\
&\qquad + \mathrm{KL}(Q(t_d|\{ct^d_c, z^d_c\})||P(t_d)) \\
&\qquad + \mathrm{KL}(Q(u^d_c|c^d_c, z^d_c)||P(u^d_c)) \\
&\qquad - \mathbb{E}_Q[\log P(X_d|t_d) + \log P(c^d_c|t_d) \\
&\qquad + \log P(X^d_{c,g}|l^d_c, z^d_c)],
\end{aligned}
$$

where KL denotes the Kullback–Leibler divergence [11]. Relative to the ELBO in scANVI, our formulation introduces three additional terms and one modification. In the reconstruction loss, there are two additional terms $\mathbb{E}_Q[\log P(X_d|t_d)]$ and $\mathbb{E}_Q[\log P(c_c^d|t_d)]$. In the KL term, we add $\mathbb{E}_Q[\log \frac{Q(t_d|c_c^d, z_c^d)}{P(t_d)}]$. In addition, instead of $\mathbb{E}_Q[\log \frac{Q(z_c^d|X_{c,g}^d)}{P(z_c^d|c_c^d)}]$, we have $\mathbb{E}_Q[\log \frac{Q(z_c^d|X_{c,g}^d)}{P(z_c^d|t_d, c_c^d)}]$. Following the adversarial training procedure in the scVI package [12], we optionally incorporate adversarial loss to improve alignment of both cell and individual embeddings across cohorts.
